# Probing the Modularity of Megasynthases by Rational Engineering of a Fatty Acid Synthase (FAS) Type I

**DOI:** 10.1101/379156

**Authors:** Alexander Rittner, Karthik S. Paithankar, David Drexler, Aaron Himmler, Martin Grininger

## Abstract

Modularity is an aspect of a decomposable system with a coordinating authority that acts as a glue which holds the loosely held components. These multi-component entities (“modules”) facilitate rewiring into different designs allowing for change. Such modular character is a fundamental property of many biological entities, especially the family of megasynthases such as polyketide synthases (PKSs). The ability of these PKSs to produce diverse product spectra is strongly coupled to their broad architectural modularity. Decoding the molecular basis of modularity, i.e. identifying the folds and domains that comprise the modules as well as understanding constrains of the assembly of modules, is of utmost importance for harnessing megasynthases for the synthesis of designer compounds. In this study, we exploit the close semblance between PKSs and animal FAS to re-engineer animal FAS to probe the modularity of the FAS/PKS family. Guided by structural and sequence information, we truncate and dissect animal FAS into its components, and reassemble them to generate new PKS-like modules as well as bimodular constructs. The novel engineered modules resemble all four common module types of PKSs and demonstrate that this approach can be a powerful tool to create higher catalytic efficiency. Our data exemplify the inherent plasticity and robustness of the overall FAS/PKS fold, and open new avenues to explore FAS-based biosynthetic pathways for custom compound design.

## Introduction

Modularity is a widespread concept found across multiple disciplines has always attracted considerable attention among industrial engineering, computer science and even natural sciences. Within this concept, individual structural elements or functions that are interdependent are clustered as modules, which in a highly independent manner form the ‘larger’ system. The complexity of such a modular system is determined by the choice of the module and the connection of such modules via few connector hubs. On the other hand, a non-modular system is characterized by optimization of all the individual units and their connectivity (see Fig. 1A).^1^ Modularity is thought to be beneficial in a changing environment as it encourages creation of variety, utilization of similarities and reduction of complexities. This leads to faster adaptation rates of the systems. The crucial aspect whether or not a non-modular system evolves a modular character seems to be a selection pressure on the minimization of connection costs alongside with maximization of the performance.^2^ A selection pressure on performance alone seems rather to lead to a fine-tuning of the existing non-modular system.^2^

**Figure 1:**
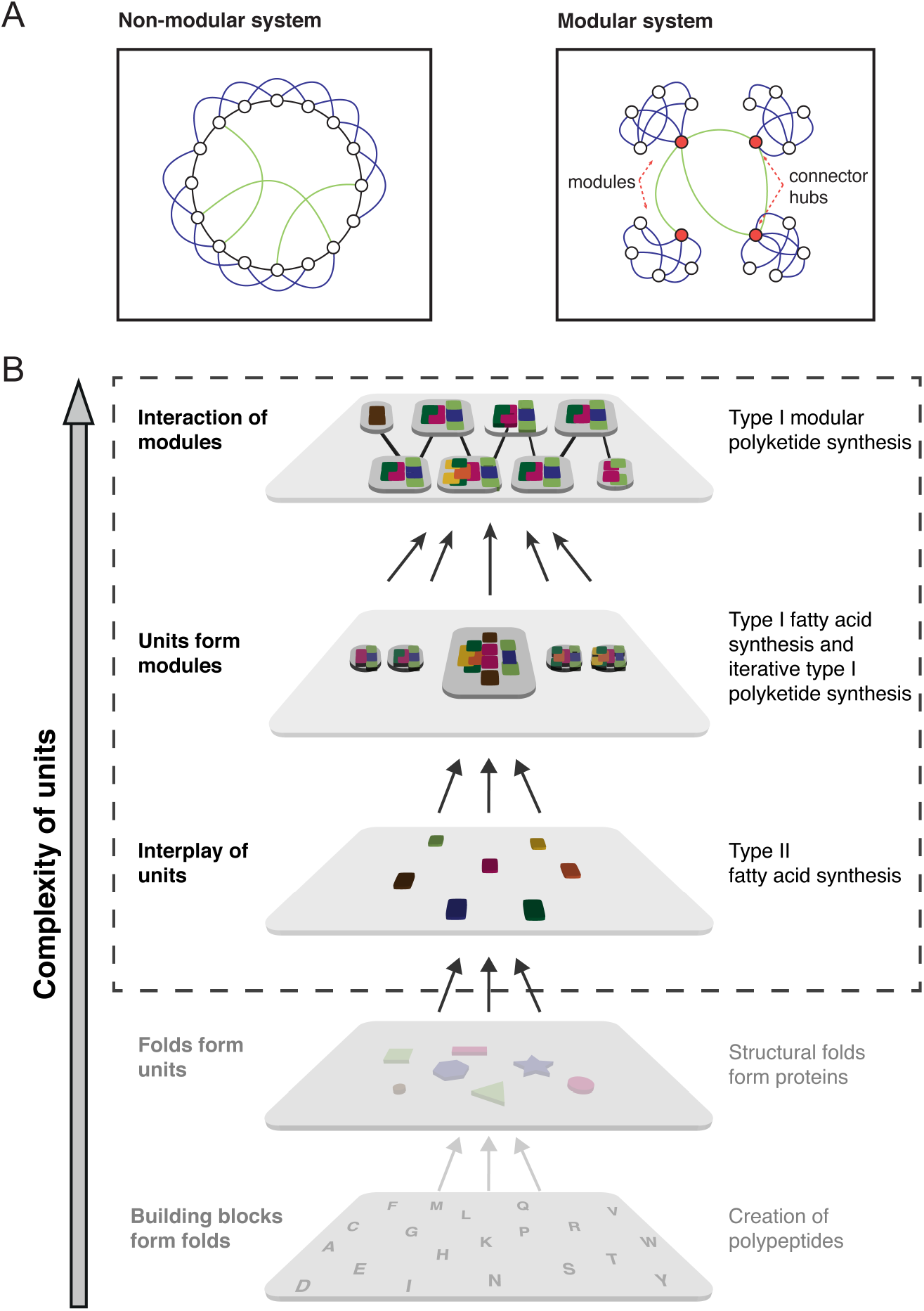
Modular vs. non-modular organization of systems. (A) Representative depiction of a modular versus non-modular organizations of a system, as described by Shanahan for the connectivity of animal brains.^1^ (B) Simplified cartoon representation of type I and type II fatty acid and polyketide synthesis. Whereas, the type II fatty acid synthesis contains major aspects of a non-modular system, modular type I PKSs have evolved to work primarily with characteristics of a modular system. A unit is defined as an enzymatic function, which structurally appears in monofunctional enzymes or domains. The depiction implies that the concept of modularity can be applied beyond our definition down to the individual amino acids as building blocks and further.

Fatty acid synthesis is an essential process in living cells that contains characteristics of non-modular as well as modular systems (see Fig. 1B). In plants, bacteria and in mitochondria, the biosynthesis is conducted by separate enzymes known as the type II synthesis. Defining an individual enzymatic function as a unit of the overall system, this reflects a fine-tuned non-modular organization embedded in the metabolism of the cell. It is tempting to speculate here that every single active site and the interactions between the separate enzymes have evolved towards a high performance in the specific task of *de novo* fatty acid synthesis. In *Corynebacterium, Mycobacterium,* and *Nocardia* (CMN) group bacteria, fungi and higher eukaryotes essentially the same enzymatic functions (see Fig. 2A) have clustered to form modules creating multidomain (type I) proteins of up to megadalton sizes. During the course of evolution, genes that lined up on genome level fused to form the type I systems, supported by the insertion of scaffolding domains that stabilize the partly elaborate architectures.^3-5^ The origin of fatty acid biosynthesis in large complexes (CMN bacteria and eukaryotes) however, is still a matter of speculation. A benefit of type I FAS mediated synthesis lies in the kinetic efficiency of the compartmentalized reaction and a coordinated expression and regulation of multidomain proteins.^6^

**Figure 2:**
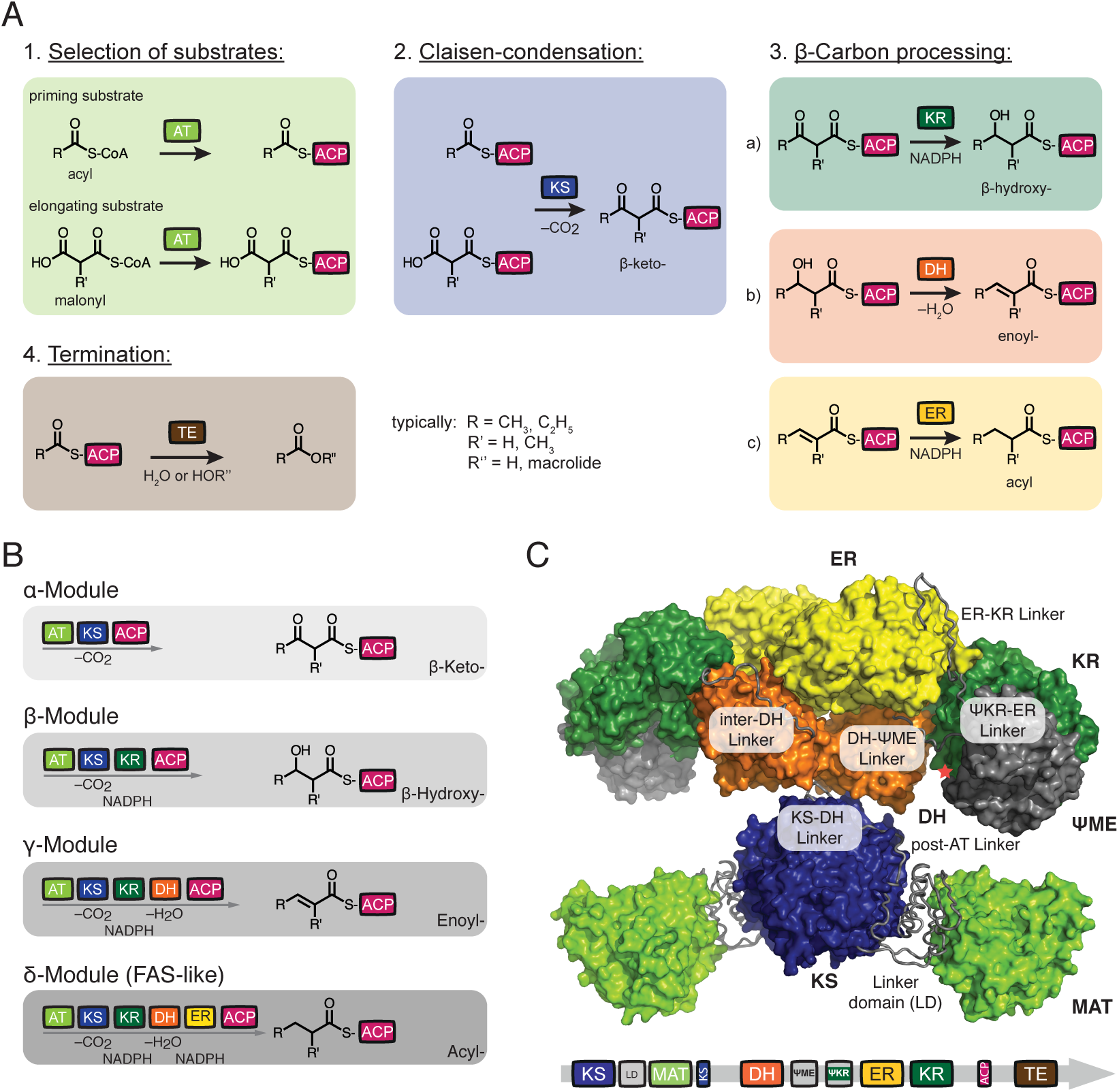
Architecture of animal FAS and PKS. (A) Commonly used domains within of FASs and PKSs. Domain nomenclature: MAT, malonyl-/acetyltransferase; AT, acyltransferase; KS, ketosynthase; KR, ketoreductase; DH, dehydratase; ER, enoylreductase; TE, thioesterase; ACP, acyl carrier protein; LD, linker domain; ψKR; truncated structural KR fold; ψME; truncated structural methyltransferase fold. (B) Domains are further connected to organize modules in FAS/PKS proteins. Essentially four module types appear in FAS/PKS (*cis*-AT PKS) family, and are termed α-(KS-AT-ACP), β-(KS-AT-KR-ACP), γ-(KS-AT-DH-KR-ACP) and δ-modules (KS-AT-DH-ER-KR-ACP). The different modules elongate and modify the polyketide intermediate. In a first of putatively several elongation cycles, R and R’ refer to typically CH_3_ and C_2_H_5_ as well as H and CH_3_, respectively. Color code as introduced in (A). (C) X-ray structure of animal FAS with domains in surface and inter-domain linkers in tube representation, respectively (porcine FAS, pdb accession code 2vz9).^14^ Red asterisk indicates the C-terminus of the KR domain, where the ACP and TE domains are flexibly tethered. TE and ACP could not be traced in electron density in the X-ray structure. Color code as introduced in (A).

The phenomenon of modularity is clearly manifested in the modular polyketide synthase (PKS) system.^7^ An optimal reuse of related modules together with variable composition of units, has led to the evolution of these class of enzymes creating compounds of vast diversity and remarkable complexity, e.g. maitotoxin.^8,9^ These polyketides are mainly produced in microorganisms to provide an advantage in a challenging environment as secondary metabolites, e.g. antibiotics, and serve as a rich source of pharmaceutically relevant chemical entities. This enzyme class shows many characteristics of a modular system as strongly interdependent units are clustered in modules (see Fig. 2B) and these modules are rather independently connected via few connector hubs. This aspect makes these enzymes highly interesting for rational protein design,^10,11^ and computational analysis suggests that employing modularity, modular PKSs can potentially produce hundred of millions of polyketide compounds.^12^

In this study, we use an animal FAS, which is evolutionary related to modules from PKSs, to investigate aspects of modularity on the protein level. Animal FAS is a suited model protein for this type of study, since it is well known in structure and function (see Fig. 2C), contains the full set of commonly used enzymatic functionalities occurring in PKSs and is catalytically highly efficient.^13-15^ By truncating and dissecting the animal FAS fold into its functional units, we probe the robustness and functional integrity of separated domains, which we then rewire to new modules and bimodular assemblies. The results of the clustering interdependent units and the presented bimodular constructs lay a clear foundation to establish a basis for FAS-based PKS-like assembly lines.^16^

## Results

### Recombinant expression of human and murine FAS

*E. coli* is a valuable heterologous host for recombinant protein production owing to low costs for cell culturing and fast and simple mutagenesis cycles. The decision to work with *E. coli* was supported by an early study on the purification of human FAS (hFAS) from recombinant expression in *E. coli*,^17^ and the previously reported successful production of the 320 kDa type I bacterial and fungal FAS in *E. coli*.^18,19^ We started to work with hFAS, because it represents a potential drug target in the treatment of cancer and obesity.^20,21^ Our attempts to access hFAS from *E. coli* expressions were either unsuccessful or yielded aggregated material, as also stated elsewhere.^22^ For detailed information on the expression of the hFAS encoding gene in *E. coli,* see Supporting Note and Figure S1 to S4. We finally also tested the production of murine FAS (mFAS) in *E. coli,* which shares 81% sequence identity to hFAS (see Fig. S5). A similar cloning strategy was conducted as performed for hFAS (see Fig. S1), resulting in a construct with a N-terminal StrepI-tag and a C-terminal His-tag. This mFAS construct was readily accessible in suited protein quality and sufficient amounts (see Fig. S4) from recombinant expressions in *E. coli*.

### Protein quality control and characterization of mFAS

In an optimized protocol, mFAS was purified by a rapid tandem affinity purification strategy (Ni-chelating and Strep-Tactin affinity chromatography), yielding 1.5-2 mg of purified protein from 1 L of *E. coli* culture (see Fig. 3A). The integrity of the protein and its oligomeric state were monitored by size exclusion chromatography (SEC) (see Fig. 3B). In agreement with earlier reports on the temperature and buffer dependent equilibrium of the monomeric and dimeric states,^23^ the main fraction of mFAS was dimeric after incubation at 37 °C, without showing significant aggregation. As reported before, *E.* coli is not capable of performing the essential post-translational modification of the ACP domain unless co-expressed with the phosphopantetheine transferase Sfp from *Bacillus subtilis*.^15^ Data showed expression of mFAS in the apo-state, and quantitative activation of mFAS in co-expression with Sfp, which was in agreement with quantitative data collected on the separate ACP. The pure, dimeric and post-translationally modified mFAS was subjected to a NADPH consumption assay with substrates malonyl- and acetyl-CoA to determine the mFAS activity (see Fig. 3C). We determined a specific malonyl consumption rate of 282 ±16 nmol min^−1^ mg protein^−1^ which was comparable to the previously reported values for recombinantly produced human FAS.^17,24^ In comparison to this, significantly higher activity (3–5 fold) have been reported for animal FASs purified from native tissues and rat FAS,^25,26^ expressed in insect cells. This compromised activity may indicate the lack of post-translational modifications, which cannot be provided by *E. coli.^27,28^* mFAS was further characterized in triacetic acid lactone (TAL) formation under non-reducing conditions in the absence of NADPH, in the tolerance to different priming substrates and by determining the steady-state kinetic parameters (see Fig. 3D and E).^29^

**Figure 3:**
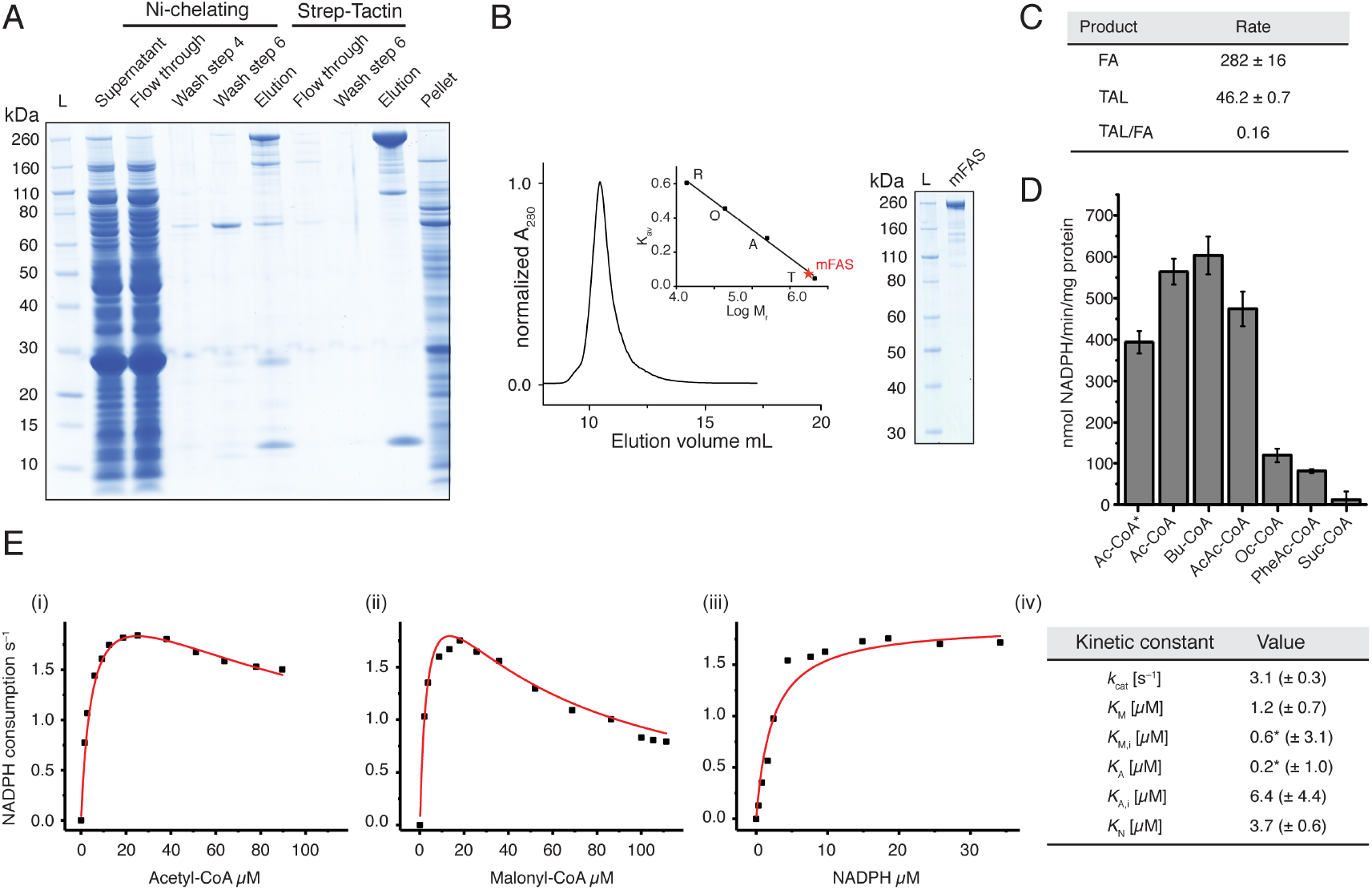
Purification and quality control of mFAS. (A) SDS-PAGE (NuPAGE 4-12 % Bis-Tris) of tandem affinity purification of mFAS, co-expressed with Sfp, using Ni-chelating and subsequent Strep-Tactin chromatography. mFAS has only little tendency to aggregate, compared to hFAS (see also Fig. S2-S4). (B) SEC of purified mFAS with absorbance normalized to the highest peak. mFAS elutes at an apparent molecular weight of 601 kDa (theoretical weight: 554 kDa). Calibration shown in inset: T, thyroglobulin; A, aldolase; O, ovalbumin; R, ribonuclease. (C) Comparison of the consumption rate of malonyl-CoA of mFAS for fatty acid (FA) and triacetic acid lactone (TAL) synthesis. The rate is given in nmol malonyl-CoA consumed per minute and mg protein. Data were collected in biological replication (n = 3). For FA synthesis, the rate of NADPH consumption was divided by two (one elongation requires two NADPH molecules) and the rate of TAL production was multiplied by two (two elongations deliver one TAL molecule), respectively. (D) Specificity of mFAS for different starter units. NADPH consumption assay was performed as described in the methods. Substrate concentrations were 100 μM X-CoA, 100 μM malonyl-CoA and 40 μM NADPH, respectively. Ac, acetyl-CoA; Bu, butyryl-CoA; AcAc, acetoacetyl-CoA; Oc, octanoyl-CoA; PheAc, phenylacetyl-CoA; Suc, succinyl-CoA. The asterisk indicates usage of a higher concentration of potassium phosphate (350 mM). Data were collected in biological replication (n = 3). (E) Typical plots of the initial velocity divided by the total enzyme concentration versus the concentration of (i) acetyl-CoA (19 μM malonyl-CoA and 30 μM NADPH), (ii) malonyl-CoA (13 μM acetyl-CoA and 30 μM NADPH) and (iii) NADPH (13 μM acetyl-CoA and 19 μM malonyl-CoA). NADPH consumption was monitored fluorometrically. For curve fitting see Methods section. The asterisk indicates that the ratio of *K*_A_/*K*_M,I_ is well defined, but the absolute value of either is not.^29^

### Deconstruction of the multi-domain architecture of mFAS

The approach of dissecting the animal FAS fold into its functional components was initiated by splitting mFAS into condensing, processing and terminating parts (see Fig. 4A). The design of the condensing KS-MAT didomain part was guided by available structural data.^14,22^ The construct comprises the native N-terminus and C-terminally ends at the waist region of mFAS. The condensing part construct expressed with typical yields of 12-15 mg purified protein from a 1 L *E. coli* culture, which corresponds to a roughly ten-fold increase in yield as compared to full-length mFAS. The processing part (see Fig. S6), starting at the waist region and extending to the C-terminal end of the KR domain, expressed at an approximately three-fold increased yield than that of the condensing part with about 38 mg purified protein from a 1 L *E. coli* culture. Expression of the domains TE and ACP, both as separate proteins, was achieved at about 100 mg and 20 mg per liter culture (see Fig. 4B), respectively. For a sequence alignment of animal FAS with domain borders highlighted, see Figure S5.

**Figure 4:**
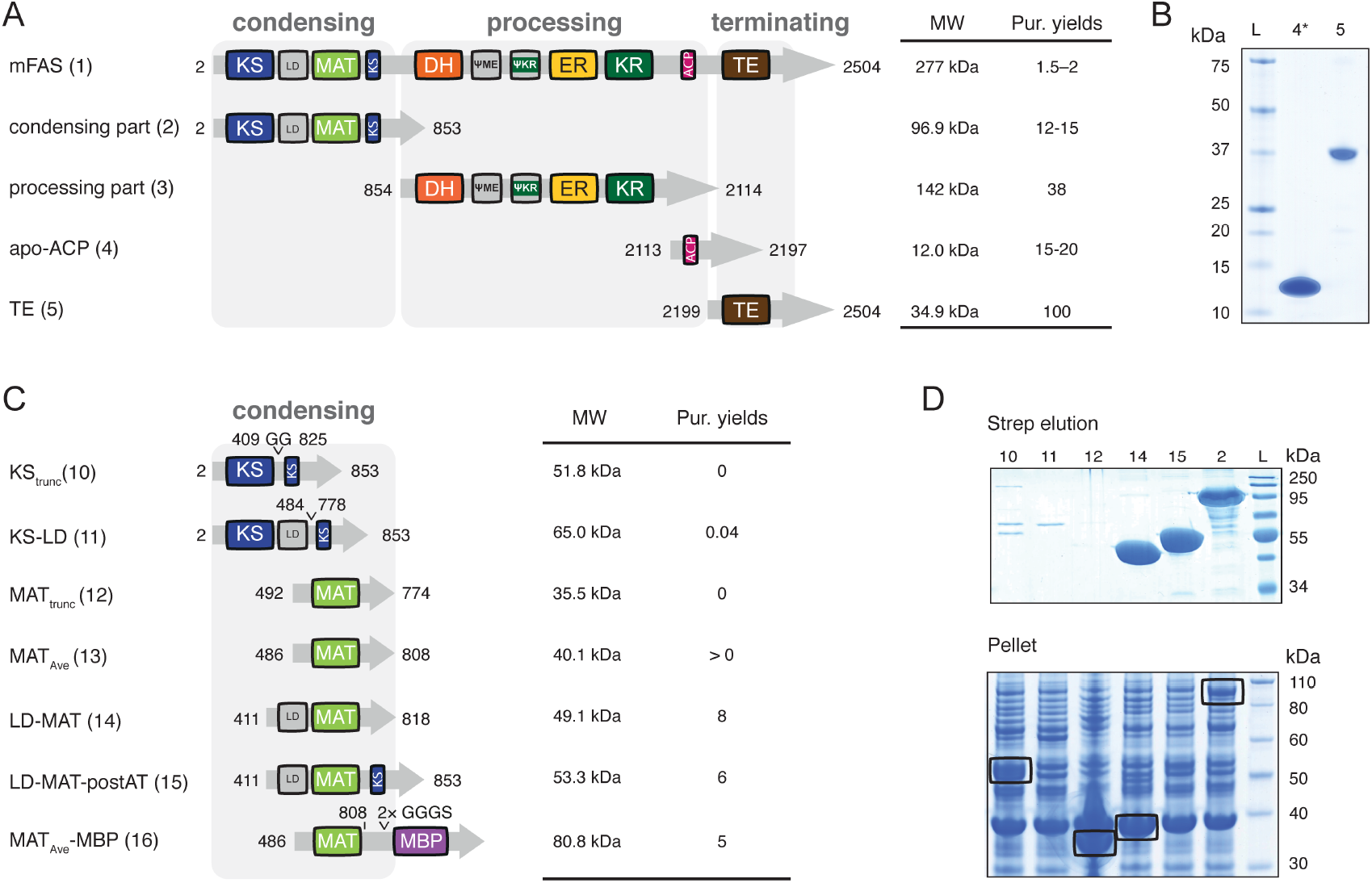
Deconstruction of the mFAS fold. All amino acid positions/numbers refer to the wild type mFAS. Molecular weights refer to proteins without the N-terminal methionine. (A) Domain organization of mFAS and first engineered mFAS constructs. Construct number are given in brackets. Constructs design was guided by sequence alignments of animal FAS (Fig. S5) and the crystal structure of porcine FAS (pdb: 2vz9).^14^ (B) Coomassie-stained SDS-PAGE gels (NuPAGE 4-12 % Bis-Tris) of constructs comprising domains TE and ACP. Star (*) indicates phosphopantheteinylated ACP received in co-expression with Sfp.^15^ (C) Domain organization of different constructs containing KS or MAT. The design is based on the crystal structures of the mFAS KS-MAT substructure (pdb: 5my0) and the avermectin loading AT (pdb: 4rl1).^15,31^ (D) Coomassie-stained SDS-PAGE gels (NuPAGE 4-12 % Bis-Tris) of constructs comprising domains of the condensing part. Aggregated protein is highlighted by black boxes. Purification yields refer to mg of protein after tandem purification from 1 L cell culture, except for ACP and TE, which were purified only via Ni-chelating chromatography.

### Engineering of the mFAS condensing part

The condensing part of the mFAS fold composed of the KS and MAT domains perform the substrate loading and condensation (see Fig 1A). Since KS-independent acyl transferases (ATs) appear in PKSs as loading ATs or as trans-ATs,^30^ we aimed at liberating the MAT from the intertwined KS-MAT fold within the mFAS (see Fig. 4C and D).

The separated MAT-domain was accessible in *E. coli* in two construct designs, which both contain the linker domain (LD), but optionally included the post-AT linker (see Fig 4C: constructs **14** and **15**) yielding about 5-10 mg of purified protein from a 1 L culture. MAT was further truncated by removing LD, based on the X-ray crystal structure of the avermectin loading AT.^31^ A minimal MAT construct (492-774; **12**) lacking a part of a β-sheet (780-783) and the long C-terminal α-helix (792-806) was not stably folded in *E. coli* as detected in the insoluble fraction (see Fig. 4D). A slightly elongated construct (486-806; **13**; termed MAT_Ave_) matching the avermectin loading AT fold was then sufficiently expressed when attaching a maltose binding protein (MBP; **16**) to the C-terminal helix. The integrity of construct **16** was confirmed by SEC and by comparable transacylation rate as for the MAT domain in the wild type mFAS as previously reported.^15^ The KS-domain in constructs with (**11**) or without LD (**10**) was hardly expressed in *E. coli* (see Fig. 4C and D).

### Engineering of PKS-like reducing Modules

Four common module types are essentially observed in modular type I PKSs (*cis*-AT PKSs); α-(KS–AT–ACP), β-(KS–AT–KR–ACP), γ-(KS–AT–DH–KR–ACP) and δ-modules (KS–AT–DH–ER–KR–ACP) (see Fig. 2B).^32^ The KR-mediated reduction of the β-ketoacyl moiety is the first step of three processing reactions that may be performed in PKSs. This KR domain is comprised of a catalytic fold and a truncated structural fold (ψKR), which is located N-terminally to the ER domain. The single crystal X-ray structure of the porcine FAS revealed that the KR domain serves as the structural hub in the processing part.^14^ It interacts with all other modifying domains, while there are few direct interactions between either of the attached domains (DH, ER and ψME) (see Fig. 5A). Whereas the same modifying domains also appear in full-reducing PKSs, the structural organization need not necessarily mimic that of mFAS. Consequently, the KR domain may not play a central structural role, as described for mycocerosic acid synthase.^33^

**Figure 5:**
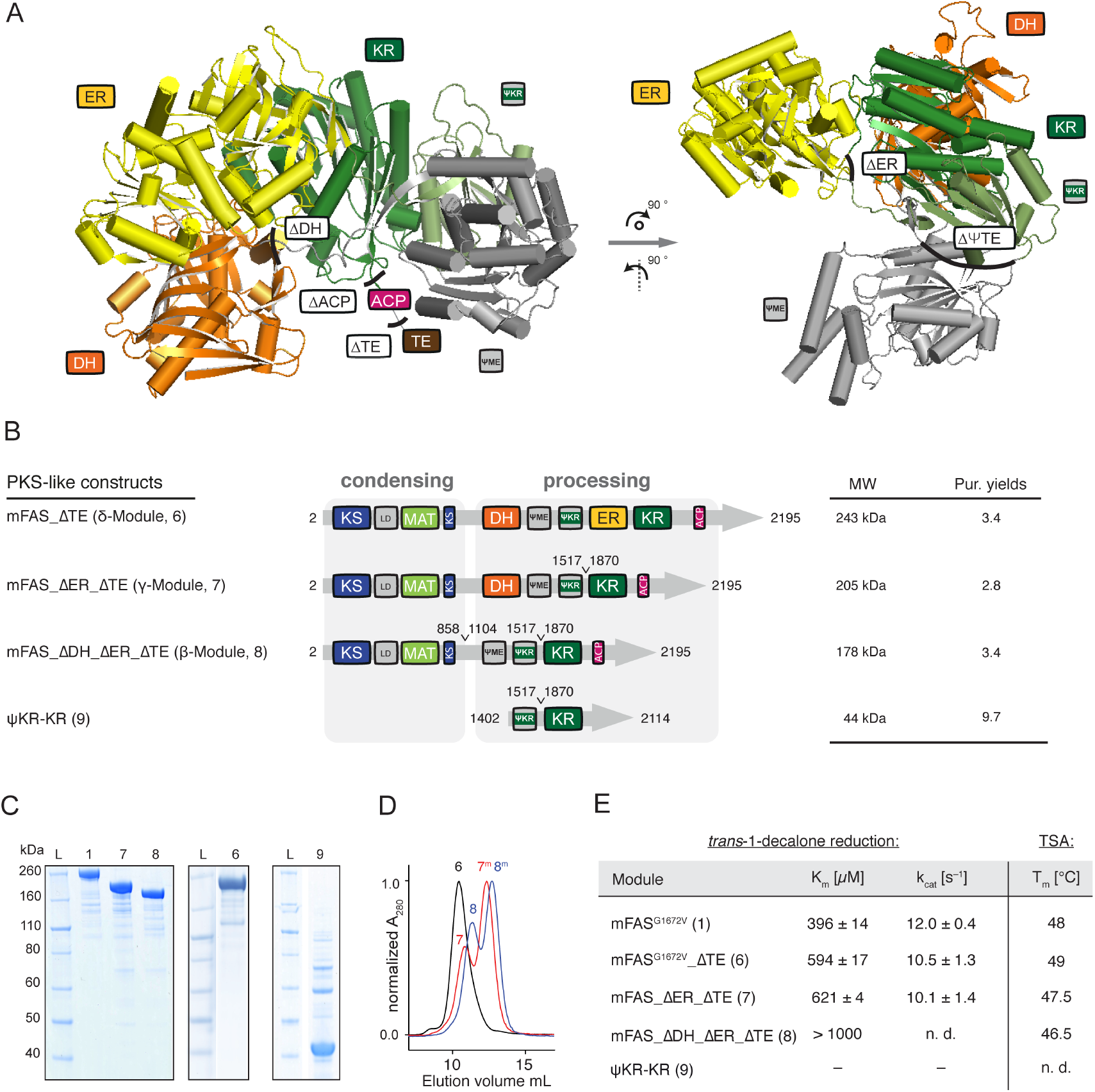
Construction of PKS-like reducing modules. (A) Structure of the monomeric processing part (porcine FAS; pdb: 2vz9) in cartoon depiction.^14^ The ψKR fold supports the integrity of the catalytic KR domain and is the docking site for the ψME domain. Black lines indicate sites used for the design on beta to delta modules. (B) Domain organization of modules rebuilding highly reducing (construct **6**; δ-module) and partially reducing (construct **7**; γ-module and construct **8**; β-module) PKSs. Purification yields refer to mg of protein after tandem purification from 1 L cell culture. (C) Coomassie-stained SDS-PAGE (NuPAGE 4-12 % Bis-Tris) of purified reducing modules (β-, γ- and δ-). Construct **9** was purified via a Strep-tactin affinity chromatography, but shows impurities. (D) SEC of purified constructs **6**, **7** and **8** with absorption normalized to the highest peak. Samples **7** and **8** were not incubated at 37 °C before SEC runs. Peaks correspond to apparent molecular weights of 584 kDa, 490 (237) and 378 (200) kDa, respectively (theoretical dimer molecular weights: **6**: 486 kDa, **7**: 410 kDa and **8**: 356 kDa). “m” refers to monomeric species. (E) Specific KR activity determined by a trans-1-decalone reduction assay and data from a thermal shift assay (TSA).^36,55^ The ER domain was knocked out by mutation G1672V;^56^ n.d. means not determined, and “–” no detectable activity. Errors refer to standard deviation from four technical replicates.

To probe the structural tolerance of the processing part, we aimed at reconstructing PKS β- to δ-modules. Construction of the δ-module (**6**; 2-2195) required the deletion of the terminating TE domain (see Fig. 5B). The TE domain is structurally independent, as observed during this study (see Fig. 4B), and from the crystal structure on the excised protein from human FAS.^34^ For construct design, the ACP-TE connecting flexible linker was preserved to attach an accessible His-tag. The construct was purified at yields of about 3 mg per liter *E. coli* culture (see Fig. 5B and C). Construct **6** was used as a template to generate the γ-module (**7**; 2-1517+1870-2195), which required the deletion of the ER domain that separates ψKR and KR and contributes to the dimeric interface of mFAS. Residues P1517 and S1870 at the N- and C-terminus of the ER domain are part of a long, accessible linker between the ψKR and KR domains and are only 5.7 Å apart from each other (distance between Cα-atoms in X-ray structural data).^14^ Bridging P1517 and S1870 extruded the ER domain, resulting in construct **7**, which expressed in good yields in *E. coli* (about 3 mg protein per liter culture; see Fig. 5B). The β-module (**8**; 2-858+1104-1517+1870-2195) was created by deleting the DH domain in construct **7**, which was achieved by linking S858 with S1104 and thereby connecting the linker of the waist region of mFAS to the full DH-ψME linker. Construct **8** expressed slightly better than construct **7** with a typical protein yield of about 3.5 mg per liter *E. coli* culture (see Fig. 5B and C). All modules were subjected to SEC, from which they eluted as a mixture of monomeric and dimeric species while showing overall little aggregation (Fig. 5D). Deletion of the ER domain pronounced the monomeric species. Protein stability was further investigated by a thermal shift assay (TSA) and melting temperatures were compared to mFAS. Interestingly, melting temperatures differed in only a few degrees implying maintained stability. Deletion of the TE domain slightly increased stability compared to that of full-length mFAS (Fig. 5E).

Besides collecting data on protein stability, we determined KR activity with the trans-1-decalone reduction assay, as a sensitive measure for the fitness of the truncated folds (see Fig. 5E).^35,36^ To exclude a potential background signal by an ER domain activity, NADPH binding sites of ER containing constructs were altered by mutation G1672V.^23^ For mFAS, kinetic constants *K*_m_ and *k*_cat_ were determined to 396 ± 14 μM and 12.0 ± 0.4 s^−1^, respectively. They compare well to values for FAS purified from pig liver of 150 μM (170 μM) and 0.73 s^−1^ (51.5 s^−1^) for both individual enantiomers 9S (9R).^35^ Construct **6** (δ-module) showed a roughly 1.5-fold increase of *K*_m_ (594 ± 17 μM) and an essentially unchanged *k*_cat_ (10.5 ± 1.3 s^−1^). The TE domain is loosely connected (owing to the flexible linker) to the processing part, and the observed increase of *K*_m_ by the TE deletion on KR function cannot be explained with current data. Construct **7** (γ-module) gave similar values for KR activity as construct **6** (*K*_m_ of 621 ± 4 μM and *k*_cat_ of 10.1 ± 1.4 s^−1^). The assay set up did not allow determining accurate values for construct **8** (β-module), but the *K*_m_ was estimated to be clearly increased as compared to the other constructs (see Fig. 5E). A stand-alone KR (**9**) only containing ψKR and KR was solubly expressed, but did not show KR activity (see Fig. 5A-C and E). See Figure S6, for engineering of the processing part as constructs separated from the condensing part.

### Engineering of a PKS-like non-reducing α-Module

Engineering of a non-reducing α-module (see Fig. 2B) requires the direct linkage of the ACP domain to the condensing part. For an initial KS-ACP linker design, sequences of KS-ACP linkers occurring in module 3 and module 11 (both non-reducing α-modules) of the avermectin synthase (AVES 2: Uniprot number Q9S0R7; AVES 4: Q79ZM6) were used as templates (see Fig. 6A). C-terminal ends of the post-AT linkers, defining the N-terminus of the designed linker, were identified by localizing a conserved motif occurring in most PKSs (_AVES_KS3: YPFQHHHYW; _AVES_KS11: YAFERERFW).^37^ The N-termini of _AVES_ACP domains were determined by homology to the ACP domain of DEBS module 2 (PDB: 2ju2), which has been determined in a structural model, and were defining the C-terminus of designed linkers.^38^ Applying this information, the selected linkers were 31 (_AVES_M3L) and 37 (_AVES_M11L) amino acid residues in length, respectively (see Fig. 6B). The linkers were inserted between the C-terminus of the KS domain and the N-terminus of the ACP domain. For identifying the N-terminus of the mFAS ACP domain, structural information of rat ACP (PDB: 2png) was considered.^39^ In the first generation of α-modules, we used the C-terminal end of the conserved motif of the post-AT linker as anchor points for linkers _AVES_M3L (**18**) and _AVES_M11L (**20**) assuming that the residual stretch of the post-AT linker only transiently interacts with the KS domain (see Fig. 6B). Furthermore, two other constructs were designed that only utilize amino acid sequences of mFAS by bridging the first β-sheet of the DH domain to the natural KR-ACP linker (linker length: 33 aa) to either connect the ACP domain (construct: **17**;) or the ACP-TE didomain (**22**). Constructs **18** and **20** yielded only about 25-30% of purified protein, as compared to the yields from **17** and **22**, although **18** and **20** did not show an increased tendency towards aggregation (see Fig. 6C). Constructs **18** and **20** yielded exclusively monomeric protein that did not dimerize, but aggregated under incubation at 37 °C. In contrast, construct **17** and **22** were purified as mixtures of monomers and dimers, which could be assembled under appropriate conditions.^23^ To determine whether the poor protein yield of constructs **18** and **20** originates from artificial avermectin linkers or the missing C-terminal stretch of the post-AT linker of mFAS, we exchanged the first 11 amino acid residues of the AVES linkers by the respective mFAS amino acid sequence to create constructs **19** and **21** (see Fig. 6A and B). These constructs showed essentially the same behavior as **17** and **22** in respect to purification yields, the ratio between monomers and dimers and the ability to assemble in high ionic strength buffers.

**Figure 6:**
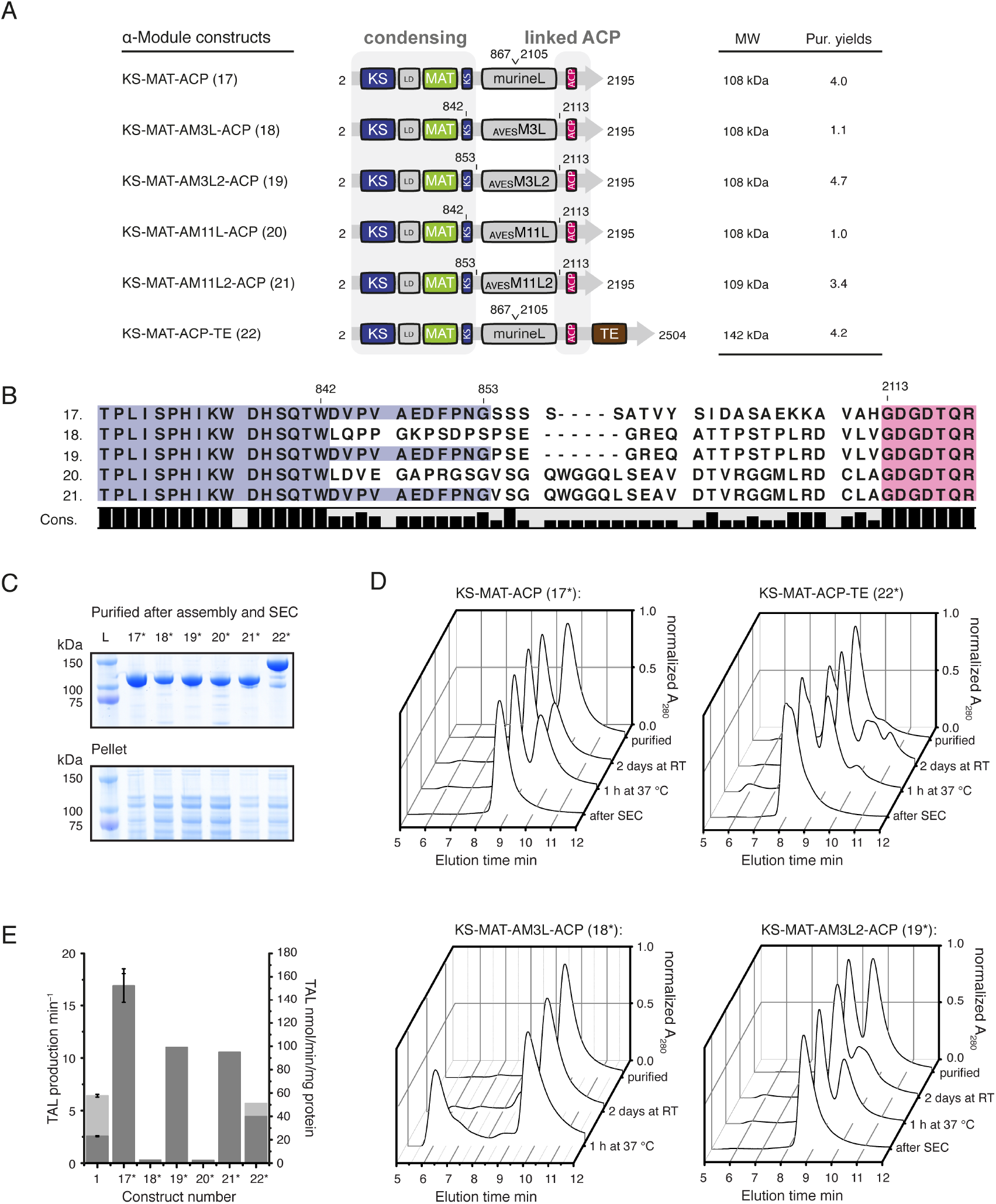
Engineering of a PKS-like non-reducing Module. (A) Domain organization of α-modules. The PKS AVES modules 3 and 11 serve as a template for linker design. (B) Sequence alignment supporting linker design. The mFAS KS domain (post-AT linker) is shown in blue and mFAS ACP domain in purple. (C) Coomassie-stained SDS-PAGE (NuPAGE 4-12 % Bis-Tris) of purified α-modules (upper panel) and insoluble fractions (lower panel). Asterisk (*) indicates phosphopantetheinylated ACP received by co-expression with Sfp (D) HPLC-SEC analysis of select constructs with absorption normalized to the highest peak. The chromatographic profiles indicate an equilibrium between the monomeric and dimeric state. For multiangle light scattering data see Fig. S7. (E) TAL assay of purified constructs after preparative SEC. Rates are given in molecules produced per minute (light grey) by dividing by the total enzyme concentration and in TAL production per minute (see Fig. 2C) and molecules produced per mg protein (dark grey). TAL production assay was performed as described in the methods section. Error bars reflect the standard deviation of biological triplets.

Constructs **17**, **19**, **21** and **22** were further purified by size exclusion chromatography to minimize contaminations with monomeric and hence inactive protein (see Fig. 6D). These samples were used to determine rates of TAL formation to provide a quantitative measure to evaluate different construct designs. Monomeric proteins **18** and **20** did not show any activity and served as negative controls. Construct **22**, with the fused ACP-TE didomain, possessed a rate of 5.7 ± 0.3 min^−1^, which was comparable to the rate of the wild type mFAS (6.4 ± 0.1 min^−1^). This reduced activity of construct **22**, may also be explained by lower protein quality (e.g. protein degradation, see Fig. 6C and D). Remarkably, constructs **17**, **19** and **21** were significantly more efficient in TAL production than mFAS, which suggests a beneficial effect of the TE truncation on the enzymatic activity of α-modules. Furthermore, also the linker to the ACP domain influences catalytic efficiency as construct **17** (16.5 ± 1.6 min^−1^) is about 50% faster than **19** (10.7 min^−1^) and **21** (10.3 min^−1^). Considering the rate per mg of protein, construct **17** (152 ± 15 nmol min^−1^ mg protein^−1^) was about 6-7 times faster than wild type mFAS (23 ± 0.5 nmol min^−1^ mg protein^−1^).

### Generation of bimodular constructs

To probe whether we can assemble functional mFAS components to new architectures, we engineered mFAS with N-terminal fusions of loading domains. These constructs mimic the N-terminal part of PKS synthetic pipelines, which comprise the priming step of polyketide synthesis as well as the first elongation step. DEBS and AVES provide a simple solution for such a design, and feature single loading didomains (AT-ACP didomain) connected to the first module (“module 1”).^31,40^ Another solution may be derived from the Pikromycin synthase (PikA), which uses a modified α-module (KS, AT and ACP) with a decarboxylating KS domain and an AT domain, specific for extender substrates.^41,42^ Both architectures were considered in our approach of engineering bimodular constructs with either mFAS sequences alone or including loading domains from modular PKSs.

To connect the ACP domain of the loading module with the KS domain of the elongation module, the natural murine ACP-TE linker was extended with three alanine residues. With this design, the linker mimics typical architectures found in modular PKSs. Construct **23** was created by linking two α-modules (construct **17**) in sequence. Constructs **24** and **25** were designed by fusing either the α-module **17** or the MAT-ACP didomain (resembling construct **14**; see Fig. 4C) to the full-length mFAS (see Fig. 7A and B). Further, loading didomains from AVES and DEBS including their natural ACP-KS linkers were N-terminally fused to full-length mFAS, generating constructs **26** and **27** (see Fig. 7A and B). Domain borders in AVES and DEBS were defined by sequence alignments, in which the N-terminus of the KS domain can be identified by sequence conservation.^43^ Finally, construct **28** resembling the architecture of AVES (module 0–1) was built solely using the domains of mFAS. A modified version of the minimal MAT fold of construct **13** (MAT_Ave_) was fused to ACP, and this loading didomain was connected to mFAS via two artificial GGS linkers at the length of the AVES linker.

**Figure 7:**
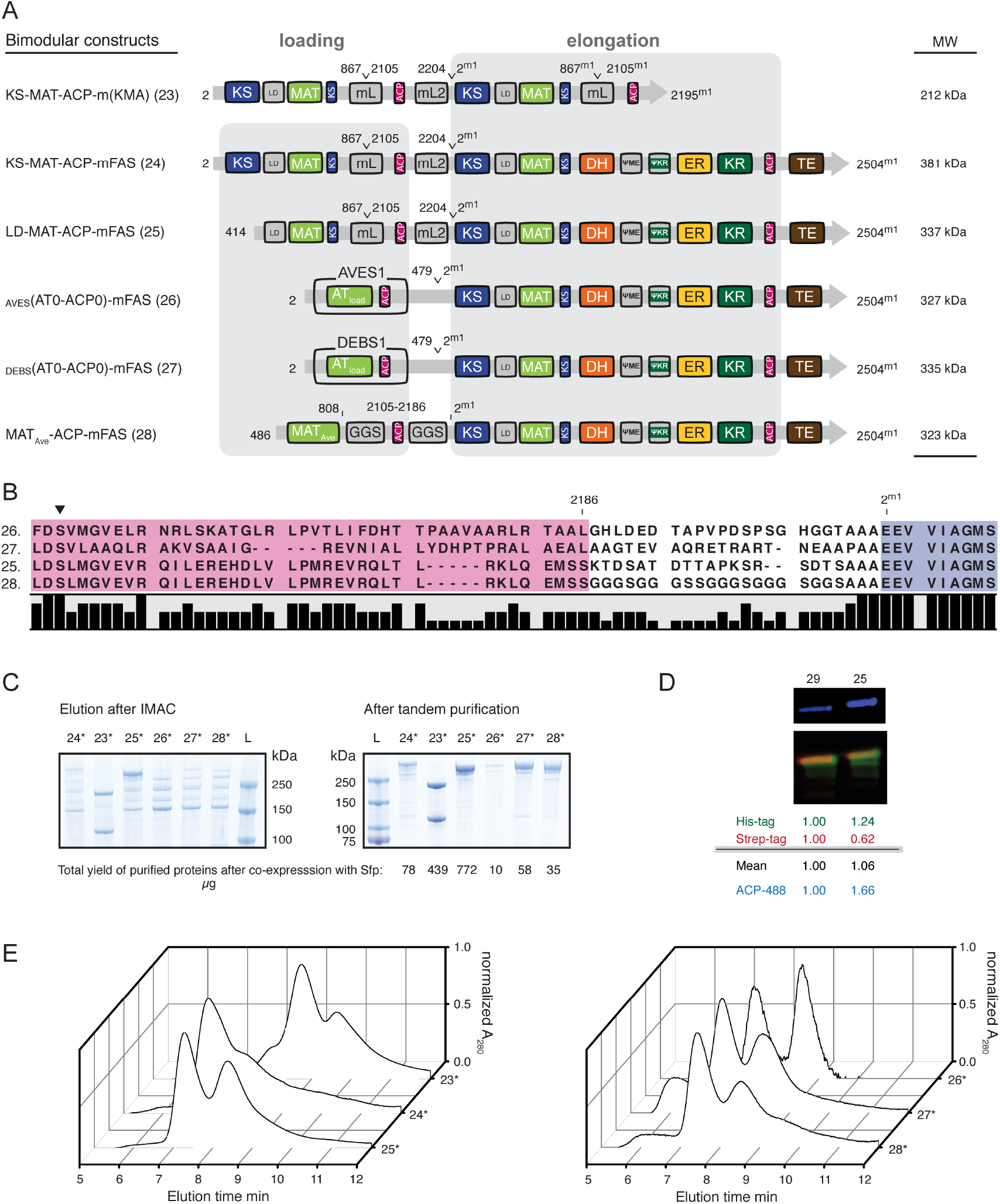
Engineering of bimodular constructs. (A) Domain organization of bimodular constructs. mFAS is used as elongation module (“module 1”) and different loading modules are fused N-terminally. Superscript “m1” indicates mFAS domain borders of “module 1”. Constructs **23-25** are fusions of mFAS with functional units of mFAS exclusively using sequences from mFAS also for linkage. Additionally, loading modules from AVES (AVES 1: Q9S0R8) and DEBS (DEBS 1: Q03131) were utilized. (B) Sequence alignment of select constructs showing sequences of the ACP to KS linker region. The mFAS KS domain is shown in blue (conserved motif: EEVVI, compare DEBS module 1: EPVAV) and the attached ACP domains in purple. The active site serine phosphopantetheinylated during post-translation modification is highlighted by a black triangle. (C) Coomassie-stained SDS-PAGE (NuPAGE 4-12 % Bis-Tris) of purified bimodular constructs after IMAC and tandem purification. Despite purification via tags at both termini, strong bands for truncated proteins remained in preparations of **23** and **24**, likely resulting from the co-purification by heterodimer formation. Protein yields are given in μg and refer to yields of purified protein (tandem purification) after coexpression with Sfp (Asterisk (*)) from 1 L cell culture. (D) Phosphopantetheinylation of ACPs in bimodular construct **25**. Quantitative Western blot after *in vitro* phosphopantetheinylation with CoA-488. A merged picture is shown in false colors. Antibodies against both tags were used. The anti-Strep antibody is colored in red and the anti-His antibody in green. Blue channel refers to fluorescently labeled ACPs. Normalized intensities for all channels are shown below in reference to control **29** (LD-MAT-mFAS). (E) HPLC-SEC of activated bimodular constructs after tandem affinity chromatography with absorption normalized to the highest peak. The two peaks refer to the monomeric and dimeric states. For multiangle light scattering data see Fig. S7.

Bimodular constructs were expressed in *E. coli* in co-expressions with Sfp, and were subsequently purified (see Fig. 7C). Constructs using natural PKS loading didomains **26** and **27** as well as constructs **24** and **28** were poorly expressed and suffered from strong proteolysis. Constructs **23** and **25** were expressed and purified in moderate yields, although construct **23** yielded heterogeneous fractions due to dimerization of a full-length with a truncated polypeptide chain. An estimate on the phosphopantetheinylation efficiency of bimodular constructs was achieved by performing the *in vitro* fluorescence assay on construct **25** (Fig. 7D). Compared to a control construct possessing only one ACP (**29**; LD-MAT-mFAS), in-gel fluorescence intensity of construct **25** was 1.66-fold increased, which agrees with the availability and accessibility of two ACP domains in this construct. HPLC-SEC showed the typical appearance of a mixture of monomers and dimers in the purified sample with an increased monomeric species compared to mFAS (Fig. 7E).

Constructs **25** and **28** appeared to be active in the NADPH consumption assay and construct **25** has also been analyzed in the interaction of the loading didomain with the C-terminal full-length mFAS.^15^

## Discussion

Here we present a rational protein engineering approach using murine FAS (mFAS) as a model protein, to study modularity as a property of megasynthases. The term modularity is generally used to describe a system, in which self-contained functional units cooperate to fulfill “greater benefits”. The variation in how units are combined to form modules and how these modules are arranged and interact within an overall architecture defines the versatility and turnover of product synthesis. A successful biotechnological application of this concept is highly dependent on the robustness and integrity of individual units, i.e. catalytic domains, and the flexibility of how these units can be reassembled with low connectivity costs to yield an overall high catalytic efficiency. While modularity has been harnessed on a gene-level, e.g. by rearranging PKS genes for combinatorial PK synthetic approaches,^44^ our goal was to assess modularity at a protein level to reveal and quantify engineering effects. Considering the evolutionarily strong connection of FASs and PKSs, some of these findings may also apply to the large and diverse family of PKSs.

Our analysis began by investigating the structural independence of individual units from the mFAS fold. A multitude of data demonstrates that individual enzymatic domains can be liberated from the structural embedment in the overall fold and remain intact. Befitting examples for that aspect are the MAT, ACP and TE domain, which can be purified as freestanding proteins that maintain their functionality.^15,34^ Furthermore, multidomain subunits as the condensing or processing part were readily accessible by truncation at interconnecting linkers, which were confirmed as suited sites for engineering.

The design of PKS-like reducing β-, γ- and δ-modules by subsequent deletion of modifying domains confirmed that also the processing part itself has a strong modular character and that the overall architecture allows relatively easy alterations. All changes were tolerated quite well yielding soluble protein with a minimal tendency to aggregate. The fitness of the folds was judged by the yield of purified protein (this measure has been used in literature before, see references),^45,46^ behavior in SEC and stability of the constructs in thermal shift assays. This suggests that individual domains in the FAS fold have maintained a high degree of independence, contrary to expectations based on the crystal structure,^14^ similarly as found in comparable iterative reducing PKSs.^33^

Nevertheless, our studies have also revealed some limitations or rather aspects that have to be considered when one intends to engineer megasynthases. A deletion of the ER domain significantly reduced the dimeric interface of PKS-like modules, which led to an increase of the monomeric protein fraction and hence inactive species. Such modules may need an optimization of remaining dimeric interfaces or, as already employed in PKSs, an introduction of stabilizing structural motifs like dimerization elements or docking domains.^47,48^ Furthermore, remaining domains although being structurally relatively independent, may be functionally impaired due to missing stabilizing effects from neighboring domains. Partial activity assays and enzyme kinetics provide a useful tool to quantify these effects and can be used to investigate specific domain-domain interactions. Hence, the activity of the KR domain in mFAS is essentially unaffected by the presence or absence of the ER domain, whereas a deletion of the DH domain dramatically reduced the catalytic efficiency, which might be explained by a stabilizing effect of α-helix 989-999 on the Rossmann fold. Moreover, deletion of the TE domain (δ-module), which is a relatively independent domain without any direct structural interface, altered the specific activity of the KR domain. This was an interesting finding as the KR domain always serves as an anchor point for the downstream domains in modular PKSs.

A further aspect of the modularity concept and a promising result for its application was demonstrated by the reassembly of the KS-MAT didomain with the ACP domain to create a non-reducing PKS-like α-module. Although the KS-domain of FAS is relatively specific for fully reduced carbon chains,^49^ it has the ability to accept β-keto groups and can condense acetyl-CoA with malonyl-CoA twice to produce triacetic acid lactone.^50^ This experiment revealed two major aspects: i) the C-terminal part of the post-AT linker, even behind the conserved motif, is crucial for the dimerization and activity of the KS domain and any alteration in its sequence has dramatic effects, at least in mFAS. This means that this part of the so-called linker should rather be seen as structural element of the KS domain than as an interdomain linker, which may differ in some modular PKSs.^51^ ii) A different design (mFAS and construct **22**) and different linkage of the identical domains KS, MAT and ACP resulted in quite remarkable differences in the catalytic efficiency to produce TAL. The simplest construct **17**, which is basically a deletion construct using exclusively mFAS sequences, possessed highest turnover rates (152 ± 15 nmol min^−1^ mg protein^−1^) demonstrating the effect of reducing connectivity costs (by minimizing complexity). Remarkably, reorganization of those domains created a 6-7 fold more efficient catalyst than the full-length mFAS, considering the different masses, and even more significantly than the iterative PKS methylsalicylic acid synthase (MSAS) from *Penicillium patulum* (12.5 nmol TAL min^−1^ mg protein^−1^).^52^

On the highest level of unit complexity, we investigated the interconnection of FAS-based modules. The simplest bimodular system was created by a N-terminal fusion of loading didomains to the mFAS fold. Remarkably, such constructs were still expressible in *E. coli*, though the yield of purified protein as well as the protein quality was dramatically decreased depending on the respective fusion. The main obstacle using such constructs seems to be *in vivo* proteolysis, which would require a sequence optimization of the mFAS gene for *E. coli* or utilization of other hosts before proceeding with more detailed functional analysis. Nonetheless our initial experiments confirmed the dimeric and potentially active state of all constructs and we could already proof a kinetic influence of the loading didomain on fatty acid synthesis in a previous study.^15^

## Conclusion and Outlook

Our study clearly demonstrates that mFAS is a modular system, which can be dissected into units that largely maintain their functionality. Furthermore, these units can be reassembled into new modules that have altered catalytic properties. The open architecture of animal FAS with few additional structural elements is well suited for engineering approaches and hence a continued development in FAS-based catalysis compared to the barrel-structured FASs.^16^

Our findings reveal opportunities and limitations of FAS engineering and provide crucial information for further applications. The core fold may readily be exploited for the synthesis of alternative products, as e.g. alcohols, by exchanging or adding other domains, e.g. exchanging the thioester domain by reductive domains.^53^ Moreover, new modules based on domains of mFAS may be used to build PKS-like assembly lines in a controlled manner with potentially high catalytic efficiency. Modularity present in this class of molecular machines may allow for creation of variety of products, by hierarchical restructuring, functional optimization by reassembling in a rational manner with minimal resources.

## Materials and Methods

pVR01 (a pET22b(+) derivative) was used as a master plasmid for cloning the human FAS gene (codon optimized; synthetic gene from Mr. Gene (GeneArt, ThermoFisher)) encoding a construct with an N-terminal StrepI and a C-terminal H(8)-tag. *FASN* genes from *Homo sapiens* and *Mus musculus* were purchased from Source BioScience (cDNA clones: IRAK110M20 and IRAVp968A0187D). Plasmids encoding chaperones from *E. coli* (TF, TF(N+C), DnaK, DnaJ, GrpE, GroEl and GroEs) were kindly provided by Prof. Hartl from the MPI Martinsried, whereas plasmids encoding DEBS 1, DEBS **2** and DEBS **3** (pBP144 and pBP130) were kindly provided by the Khosla laboratory at Stanford University. Microbial genomic DNA was purchased from DSMZ: *Pyrococcus furiosus* (DSM3638) and *Streptomyces avermitilis* (DSM46492). We performed PCR based cloning for deletions or point mutation and ligation independent cloning with the In-Fusion HD Cloning Kit (Clonetech) as described previously.^15^ All plasmids encoding the presented constructs are listed in the Supplemental Info (see, Table S1 Primers and information about cloning are summarized in Table S2. The sequences of all used plasmids were confirmed with the “dye terminator” method. All CoA-esters were purchased from Sigma-Aldrich.

### Expression of FAS and truncated constructs thereof

All plasmids were transformed into chemically competent *E. coli* BL21gold(DE3) cells. The transformants were grown overnight at 37 °C in 20 mL LB (100 μg mL^−1^ ampicillin (amp) and 1 % (w/v) glucose) medium. Precultures were used to inoculate 1 L TB medium (100 μg mL^−1^ amp). Cultures were grown at 37 °C until they reached an optical density (OD_600_) of 0.5–0.6. After cooling at 4 °C for 20 min, cultures were induced with 0.25 mM IPTG, and grown for additional 14–16 h at 20 °C and 180 rpm. Cells were harvested by centrifugation (4,000 rcf for 20 min). The cell pellets were resuspended in lysis buffer 1 (50 mM sodium phosphate, 200 mM NaCl, 10 % (v/v) glycerol, 1 mM EDTA, 1 mM DTT, 20 mM imidazole (pH 7.6)) or lysis buffer 2 (50 mM sodium phosphate, 450 mM NaCl, 20 % (v/v) glycerol, 1 mM DTT, (pH 7.6)) and lysed by French press. After centrifugation at 50,000 rcf for 30 min, the supernatant was mixed with 1 M MgCl2 to a final concentration of 2 mM. The cytosol was transferred to Ni-NTA-columns and washed with 5 column volumes (CV) wash buffer (lysis buffer without EDTA). Bound protein was eluted with 2.5 CV elution buffer (50 mM sodium phosphate, 200 mM NaCl, 10 % (v/v) glycerol, 1 mM DTT, 300 mM imidazole (pH 7.6). The eluent was transferred to Strep-Tactin columns, and washed with 5 CV strep-wash buffer (250 mM potassium phosphate, 10 % (v/v) glycerol, 1 mM EDTA, 1 mM DTT (pH 7.6)). Proteins were eluted with 2.5 CV elution buffer (strep-wash buffer containing 2.5 mM D-desthiobiotin). After concentration to 5–20 mg mL^−1^, the proteins were frozen in liquid nitrogen and stored at –80 °C. Samples were thawed at 37 °C for 30 min and further polished by size-exclusion chromatography (SEC) using a Superdex 200 GL10/300 column equilibrated with the strep-wash buffer. Proteins were concentrated again to 5–20 mg mL^−1^ and stored frozen in aliquots using liquid nitrogen.

Protein concentrations were calculated from the absorbance at 280 nm, which was recorded on a NanoDrop 2000c ((Thermo Fisher Scientific)). Extinction coefficients were calculated from the primary sequence without *N*-formylmethionine with CLC Main workbench (Qiagen).

### HPLC-SEC analysis

HPLC analysis of proteins was performed using a Dionex UltiMate 3000 RSLC equipped with a RS fluorescence detector. Chromatographic separation was performed on a SEC column (bioZen 1.8 μm SEC-3) in the buffer (150 mM KPi, pH 6.8) at room temperature. Proteins were detected by monitoring the absorbance at 280 nm. Dimeric FAS was found to elute at 7.5-7.6 min.

The signal for multiangle light scattering (MALS) was monitored using a miniDAWN TREOS MALS detector (Wyatt Technology Corporation) operated with a linear polarized laser light at 658 nm. Data were analyzed with the software ASTRA (version 7.1.0.29, Wyatt Technology Corporation, USA).

### Overall Fatty Acid Synthase Activity

FAS activity was measured fluorometrically with a microplate reader (CLARIOstar, BMG LABTECH GmbH) by following the oxidation of NADPH at 25 °C in 50 mM potassium phosphate (pH 7.0), 75-100 μM acetyl-CoA or other priming substrates, 100 μM malonyl-CoA and 40 μM NADPH. Alternatively, the absorbance at 340 nm was observed with a NanoDrop in cuvette mode using an extinction coefficient for NADPH of 6220 M^−1^cm^−1^. The enzyme was prepared in a 4-fold stock containing BSA (1.2 mg/mL) for stabilization, resulting in a final protein concentration (one polypeptide chain) of 20 nM and BSA (0.03 mg/mL) in the assay. The reaction was initiated by the addition of malonyl-CoA. Every measurement was performed in technical triplicates and the corresponding background (without CoA-esters) was subtracted.

Microplate reader settings were as follows: excitation: 348-20 nm; emission: 476-20 nm; gain: 1490; focal height: 5.7 mm; flashes: 17; orbital averaging: 4 mm.

### Analysis of Kinetic Data from Overall Fatty Acid Synthesis

Data was analyzed according to the methods of Cox and Hammes using the software OriginPro 8.5 (OriginLab, USA).^29^ To facilitate a global fit of the three data sets (varied substrate concentrations of NADPH, acetyl-CoA and malonyl-CoA) in Origin, the equation:

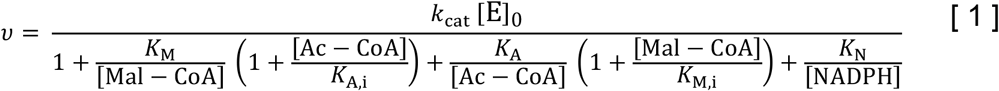

had to be converted into:

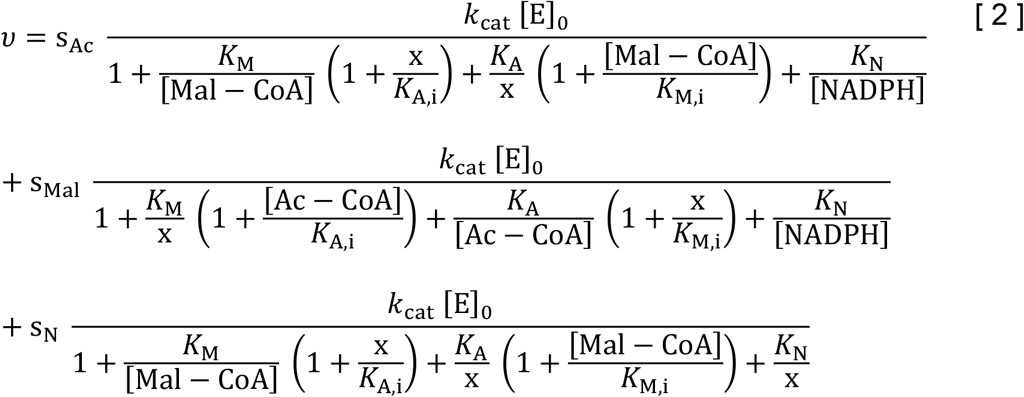

where parameters s_Ac_, s_Mal_ and s_N_ were put to zero or one assigning the summand to the respective data set. As other substrate concentrations were kept constant, all concentrations reduced to simple numbers. All other kinetic constants are defined as described in Cox and Hammes.^29^

### Triacetic Acid Lactone (TAL) Production Assay

TAL production was monitored photometrically with a NanoDrop (cuvette mode) by observing the increase in absorbance at 298 nm. Two solutions were prepared in the assay buffer (50 mM potassium phosphate, pH 7.0, 10 *%* (v/v) glycerine, 1 mM DTT and 0.03-0.05 mg/mL BSA). Solution 1 contained the substrates acetyl- and malonyl-CoA (both 200 μM) and solution 2 contained the enzyme (1400 nM) in 2-fold concentrated stock solutions. 50 μL of both solutions (incubated at 25 °C) were mixed to final concentrations of 100 μM acetyl-CoA, 100 μM malonyl-CoA and 700 nM enzyme and the absorbance was monitored for 5 min. Absorbance was converted into concentrations by using the extinction coefficient for TAL of ε = 2540 M^−1^ cm^−1^.^54^

### Specific Ketoreductase activity

Ketoreductase activity was measured fluorometrically by monitoring the consumption of NADPH at 25 °C according to previous reports.^25,35^ Two solutions were prepared in the assay buffer (100 mM potassium phosphate, pH 7.0) with solution 1 containing the enzyme and trans-1-decalone and solution 2 containing the NADPH, which was used to start the reaction. 50 μL of both solutions (incubated at 25 °C) were mixed to final concentrations of 0.05-0.15 μM enzyme, 0.02-10 mM trans-1-decalone and 300 μM NADPH and the consumption was monitored for 15 min. Fluorescence was converted into concentrations by using a calibration curve with NADPH in the respective buffer.

Microplate reader settings (Tecan Infinite M200) were: excitation: 360-20 nm; emission: 455-20 nm; gain: 110.

## Supplementary Material

Additional supporting information may be found in the online version of this article.

## Authors contribution

A.R. performed molecular cloning, protein expression, purification experiments, enzymatic assays and analyzed corresponding data. A.R. conceived the project. M.G. designed the research. D.D. established and performed specific KR kinetic experiments and the TSA with PKS-like modules under supervision of A.R.. A.H. established and performed kinetic experiments on the whole mFAS under supervision of A.R.. The authors A.R., K.S.P. and M.G. analyzed data and wrote the manuscript.

## Acknowledgments

This work was supported by a Lichtenberg grant of the Volkswagen Foundation to M.G. (grant number 85701). Further support was received from the LOEWE program (Landes-Offensive zur Entwicklung wissenschaftlich-ökonomischer Exzellenz) of the state of Hesse conducted within the framework of the MegaSyn Research Cluster.

## Abbreviations

FAS: fatty acid synthase
PKS: polyketide synthases
CMN: *Corynebacteria, Mycobacteria* and *Nocardia*
MAT: malonyl-/acetyltransferase
AT: acyltransferase
KS: ketosynthase
KR: ketoreductase
DH: dehydratase
ER: enoylreductase
TE: thioesterase
ACP: acyl carrier protein
LD: linker domain
ψKR: truncated structural KR fold
ψME: truncated structural methyltransferase fold

